# Cooperative action of separate interaction domains promotes high-affinity DNA binding of *Arabidopsis thaliana* ARF transcription factors

**DOI:** 10.1101/2022.11.16.516730

**Authors:** Mattia Fontana, Mark Roosjen, Isidro Crespo García, Willy van den Berg, Marc Malfois, Roeland Boer, Dolf Weijers, Johannes Hohlbein

## Abstract

The signaling molecule auxin is pivotal in coordinating many growth and development processes in plants mainly through the modulation of gene expression. The transcriptional response to auxin is mediated by the family of auxin response factors (ARF). Monomers of this family recognize a DNA motif (TGTC[TC]/[GG]) called the auxin-response element (AuxRE). ARFs can homodimerize through their DNA binding domains (DBD) thereby enabling cooperative binding for a bipartite inverted AuxRE (IR7). In addition to the DBD, most ARFs contain a C-terminal Phox and Bem1p (PB1) domain both capable of homotypic interactions, and mediating interactions with Aux/IAA repressors. Given the dual role of the PB1 domain, and the ability of both DBD and PB1 domain to mediate dimerization, a key question is how each of these domains contributes to conferring DNA-binding specificity and affinity. So far, ARF-ARF and ARF-DNA interactions have mostly been approached using qualitative methods that do not provide a quantitative and dynamic view on the binding equilibria. Here, we utilize a DNA binding assay based on single-molecule Förster resonance energy transfer (smFRET) to study the affinity and kinetics of the interaction of several *Arabidopsis thaliana* ARFs with an IR7 AuxRE. We show that both DBD and PB1 domains of AtARF2 contribute toward DNA binding, and we identify ARF dimer stability as a key parameter in defining affinity and kinetics seen for the DBDs of different AtARFs. Lastly, we derived an analytical solution for a four-state cyclic model that explains both the kinetics and the affinity of the interaction between AtARF2 and IR7. Our work demonstrates that the affinity of ARFs towards composite DNA response elements can be tuned by small changes of their dimerization equilibrium suggesting that this effect has major implications for ARF-mediated transcriptional activity.

## Introduction

The plant signaling molecule auxin plays a major role in many cellular and developmental processes. Auxin triggers both non-transcriptional and transcriptional responses with the latter being controlled by the nuclear auxin pathway(1–5). This pathway involves three main players: the transcription factor ARF, its repressor Aux/IAA and the ubiquitin ligase complex SCF^TIR1/AFB^. Binding of auxin to TIR1/AFB enables the recognition and ubiquitination of Aux/IAA. Upon degradation of Aux/IAA, ARF is able to modulate the expression of its downstream target genes.

The interaction between ARFs and Aux/IAAs is mediated by the C-terminal Phox and Bem1p (PB1) domain present in both proteins. The PB1 domain features two oppositely charged surfaces (type I/II or AB [acid basic] PB1 domain) that can undergo head to tail oligomerization(6–9). Remarkably, this structural characteristic enables scenarios of homo- and hetero-oligomerization among and between Aux/IAAs and ARFs. In addition to the PB1 domain, ARFs consist of two other domains, the Middle Region (MR) and the N-terminal DNA Binding Domain (DBD). The MR domain is predicted to be intrinsically disordered(5) and its amino acid sequence differs between the three phylogenetically separated ARF clades (A,B and C)(10, 11). When tested for their effect on gene expression, some ARFs activate auxin-responsive genes while other repress them. In general, class A ARFs (e.g., *A. thaliana* ARF5) act as activators while class B (e.g., *A. thaliana* ARF1 and 2) and C ARFs act as repressor(10). The DBD domain physically interacts with its DNA response element called AuxRE (auxin-responsive element)(12). This cis-regulatory element was first identified in promoters of an auxin-responsive genes in pea(13) and soybean(14, 15) and was found to be essential for their auxin inducibility. The canonical TGTCTC recognition sequence was later shown to be bound by different members of the ARF family(12, 16). More recently, the TGTCGG recognition sequence was found to have an even higher affinity for ARFs *in vitro*(17–19) and was used to create an enhanced artificial auxin response reporter(20). Single AuxREs are bound by single ARF monomers but ARF DBDs can dimerize in solution and bind cooperatively to composite response elements bearing two AuxRE in inverted configuration (IR)(17); moreover, ARF dimerization through its DBD is necessary for ARF function *in vivo*(17, 21). Interestingly, the PB1 domain seems to have diverse effects on different class A ARFs as its deletion in *M. polymorpha* ARF1 generates a loss-of-function mutant(22) whereas in *A. thaliana* ARF5 the mutant maintains its function and is hyperactive(23). The effect of the homotypic interaction of ARF PB1 domains of another class A ARF, AtARF19, has been studied using synthetic auxin response circuits in yeast, showing that mutating either the positive or the negative side of the PB1 domain reduces its ability to promote transcription(24).

Although many structures and relevant interactions among the various components of the auxin nuclear pathway have been identified, quantitative data on the affinity and kinetics of these interactions have remained scarce. In particular, the effects of the dimer/monomer equilibrium on the interaction between ARFs and between ARFs and AuxREs, or the effect of mutations on the DBD and PB1 domains on ARF dimerization have not yet been systematically studied, obscuring which interactions might be relevant in a cellular context. Particularly, it is unclear if and how both interaction domains (DBD and PB1) contribute to DNA binding, and what their relative contributions are. Furthermore, it is unclear whether oligomerization of ARF PB1 domains contributes to DNA binding.

Here, we employed a DNA-binding assay based on single-molecule Förster resonance energy transfer (smFRET) to quantitatively assess the binding affinities between different *A. thaliana* ARFs and a response element composed of two AuxREs in an inverted repeat configuration with a spacing of 7 base pairs (IR7). We found that, while the DNA-binding domain alone can bind DNA, the presence of the PB1 domain increases the affinity of AtARF2 towards the tested composite response element. In fact, this effect can be ascribed to increased stability of the dimer, whereas AtARF2 oligomerization has no sizable effect. We introduce a general four-state cyclic model to quantify the mechanisms of ARF interaction with the bipartite DNA response element; the simultaneous analysis of the equilibrium and kinetics data using this model revealed that the increase in affinity can be completely pinned to the shift in the dimer-monomer equilibrium. Further analysis of variants of AtARF5-DBD and other AtARF-DBDs showed that changes in dimer stability generated by changes in the DBD domain displays the same pattern on the kinetics as the ones generated by changes in the PB1 domain, highlighting that stable protein dimers ensure high-affinity DNA binding, no matter the source of their stability.

## Materials and Methods

### Protein expression and purification

Protein expression and purification was carried out as described previously(17). Briefly, the genomic regions corresponding to the DNA binding domain (DBD) of *Arabidopsis thaliana* ARF1, ARF2, ARF5 and full-length ARF2 were amplified and cloned in an modified expression vector pTWIN1 (New England Biolabs) to generate fusions with the Chitin Binding Domain (CBD) and Intein. ARF-CBD fusion proteins were expressed in *E. coli* strain Rosetta DE3 (Novagen). Cells were inoculated in Difco Terrific Broth (BD), supplemented with ampicillin and grown to an OD_600_ of 0.5 to 0.7, protein expression was induced by adding IPTG and the temperature was switched from 37 °C to 20 °C; the growth was continued for 20 h. Cells were harvested by centrifugation and resuspended in 50 mL extraction buffer (20 mM Tris, 500 mM NaCl, 1 mM EDTA, 0.1 % NP-40 and 2 mM MgCl_2_, pH 7.8, 10 mg of DNase and 0.2 mM PMSF). Cells were then lysed by passing the suspension twice through a French Pressure cell press and cell-free extract was generated by centrifugation. The supernatant was loaded onto a chitin column (New England Biolabs) and washed with 10 column volumes washing buffer (20 mM Tris, 500 mM NaCl, pH 7.8) using an AKTA explorer 100 (GE Healthcare). ARF-DBD proteins were eluted by 1 h incubation with 40 mM DTT in washing buffer. Proteins were concentrated using Amicon ultra-15 10K spin filters, and next passed over a Superdex 200PG size-exclusion chromatography column. ARF-DBD proteins were eluted using washing buffer with 1 mM DTT, concentrated using Amicon ultra-15 10K spin filters and stored until use at −80 °C.

### DNA constructs

Single strand DNA oligonucleotides were ordered from Eurogentec. Each strand contained a 5-C6-amino-dT modification at the desired position for labelling. Some of the strands were purchased biotinylated at their 5’end to allow for surface immobilization using a Neutravidin bridge. Strands were labelled with the desired dye (Cy3 or ATTO 647N NHS-ester) following a modified version of the protocol provided by the dye manufacturer and purified using polyacrylamide gel electrophoresis (20 % Acrylamide). DNA constructs were annealed by heating complementary single strands to 95 °C in annealing buffer (250 mM NaCl, 10 mM Tris HCl pH 8, 1 mM EDTA) followed by cooling down to room temperature overnight.

### Single-molecule FRET

Imaging was carried out on a homebuilt TIRF microscope, described previously(25). The measurements were performed using alternating-laser excitation (ALEX)(26); in this excitation scheme, each frame during which the donor is excited is followed by a frame in which the acceptor is directly excited. The emission of the fluorophores is spectrally divided into two different detection channels on the emCCD camera sensor (Andor iXon 897 Ultra). This approach creates four photon streams, three of which are relevant; (1) donor emission after donor excitation (*DD*), (2) acceptor emission after donor excitation (*DA*, arising from FRET) and (3) acceptor emission after acceptor excitation (*AA*). The three photon streams can be used to calculate the raw FRET efficiency (*E*^*^ = *DA/*(*DD* + *DA*)) and stoichiometry (*S* = (*DD* + *DA*)*/*(*DD* + *DA* + *AA*))(27). *E*^*^ contains the information about the relative distance of the two fluorophores whereas *S* contains information about the photophysical state of a given molecule (allowing to filter out molecules missing an active donor or an active acceptor). The camera acquisition time and the excitation time were set to 250 ms per frame; laser powers were set to 3 mW for green (*λ* = 561 nm) and 0.5 mW for red (*λ* = 638 nm) lasers. The PBS-based imaging buffer (pH 7.4) contained 137 mM NaCl, 2.7 mM KCl, 10 mM phosphate, and 1 mM Trolox, 1 % gloxy and 1 % glucose to decrease the rate of photobleaching(28, 29).

### Single-molecule titration experiments

Labelled dsDNA oligos were immobilized on a PEGylated glass coverslip as described previously(30). In particular, the PEGylation was carried out inside the wells of silicone gaskets placed on the coverslip (Grace Bio-labs). Each protein titration was performed using a single well, washing it between data points with 600 µL of 1x PBS buffer. The final washing step consisted of three washings separated by 15 minutes. Typically, each data point consisted of four movies (1000 frames each).

### Binding isotherms analysis

The fit of the FRET efficiency distribution with the two Gaussian distributions pertaining to the free and ARF-bound DNA populations returns an uncorrected fraction of ARF-bound DNA for each tested protein concentration 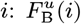. Even when no protein is added, the double Gaussian fit returns an uncorrected fraction bound 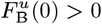 (typically ≈ 0.1). This value is an indication of the error connected to the two-population fit and can be used to renormalize the entire titration under the assumption that, in case the DNA would be completely bound by ARF, the expected uncorrected free fraction would have the same value 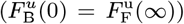. Then, the corrected fraction bound for each data point can be calculated as

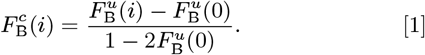

The corrected fraction bound (henceforth *F*_B_) can be fitted with the appropriate mathematical model for the interaction.

### Time traces analysi

First, the time traces from individual DNA molecules were filtered to remove sections in which either the donor or the acceptor were inactive due to fluorophore bleaching or blinking. Each molecule was allowed to take values of *E*^*^ and *S* outside the thresholds (typically 0 to 0.85 for *E*^*^ and 0.5 to 0.9 for *S*) for a maximum of three consecutive data points; longer stays outside the thresholding range resulted in the trace being interrupted. In case the molecule reentered the allowed range for *E*^*^ and *S*, the data points were saved as a new trace. The minimum length of traces was set to 50 data points (100 frames). The filtered time traces were then loaded in the software package ebFRET to perform and empirical Bayesian Hidden Markov Modelling(31). The analysis was performed assuming two states, with two restarts and a convergence threshold of 10^−6^. The results of the analysis were exported as ‘.csv’ text files and the transition matrix was used to calculate *k*_on_ and *k*_off_. The *K*_d_ was calculated as *k*_off_*/k*_on_.

### SAXS

Different concentrations of AtARF1 and AtARF5 ranging from 17 µM to 170 µM (0.7 mg mL^−1^ to 7 mg mL^−1^) were tested to record ARF dimerization depending on protein concentration. All the samples were prepared in a final buffer consisting of 20 mM Tris-HCl pH 8.0, 150 mM NaCl, 1 mM DTT. SAXS data was collected at NCD-SWEET beamline (BL11, ALBA Synchrotron, Barcelona)(32, 33). The buffer was collected for subtraction of protein samples. Measurements were carried out at 293 K in a quartz capillary of 1.5 mm outer diameter and 0.01 mm wall thickness. The data (20 frames with an exposure time of 0.5 sec/frame) was recorded using a Pilatus 1M detector (Dectris, Switzerland) at a sample-detector distance of 2.56 m and a wavelength of 1.0 Å.

Buffer subtraction and extrapolation to infinite dilution were performed by using the program package primus/qt from the ATSAS 2.8.4 software suite(34). The forward scattering *I*(0) and the radius of gyration (*R*_g_) were evaluated by the Guinier approximation, and the maximum distance *D*_max_ of the particle was also computed from the entire scattering patterns with AutoGNOM. The excluded volume *V*_p_ of the particle was computed from the Porod invariant. The scattering from the crystallographic models was computed with CRYSOL(35). The volume fractions of the oligomers were determined with OLIGOMER(36), using as probe the available PDB structures.

## Results

### The AtARF2 PB1 domain promotes DNA binding through stabilization of the AtARF2 dimer

A ChIP experiment on *Arabidopsis Thaliana* ARF19 expressed in yeast(24) suggested that PB1 mutations affect DNA-binding affinity. However, other yeast proteins may confound the differences observed in this assay. We therefore focused on the minimal system of the purified ARF protein and its DNA target *in vitro* and asked whether the interactions between the PB1 domains modulate affinity of ARFs towards a composite AuxRE. We designed smFRET experiments in which the binding of ARFs to a small doubly labelled dsDNA oligo containing two AuxREs in an inverted configuration spaced by seven base pairs (IR7) leads to a decrease of FRET efficiency (Fig. 1a, Supporting Information Fig. S1). We then performed titrations with increasing concentrations of different ARF2 variants (Fig. 1b). The ARF DBD alone is sufficient for cooperative DNA binding to the IR7 element(17); to explore the influence of regions outside the DBD, we first compared binding of AtARF2-DBD and full-length AtARF2 (FL; DBD-MR-PB1). The FRET efficiency distributions of the DNA sensor show the free DNA population (93 % occupancy) centered at *E*^*^ = 0.59 in absence of ARF proteins. With increasing concentration of AtARF2-DBD or AtARF2-FL, the low FRET population representing the ARF-bound DNA fraction (centered at *E*^*^ = 0.42) becomes progressively more populated.

**Fig. 1.**
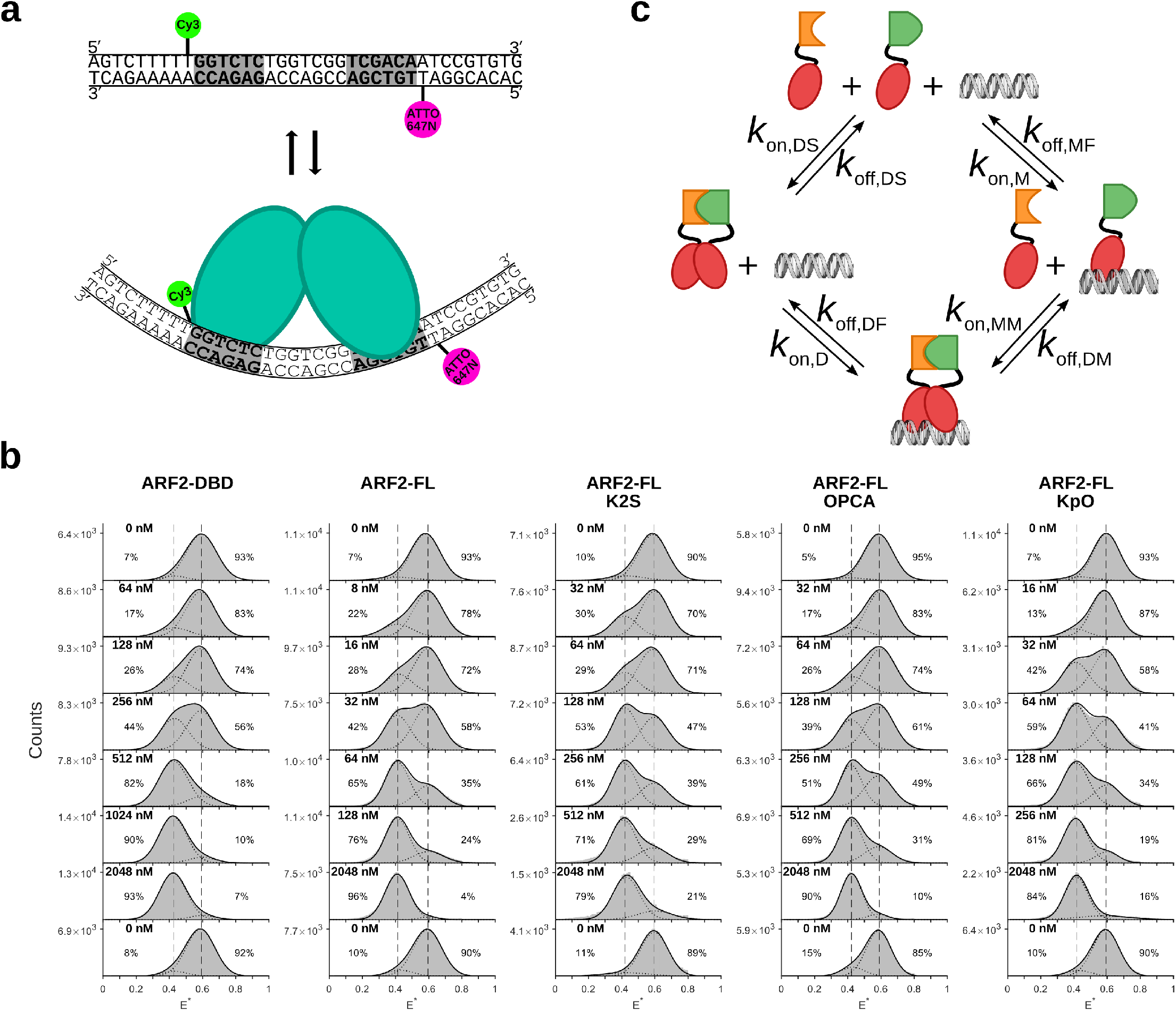
SmFRET binding assay and cyclic four-state model. (a) Schematic representation of the DNA-binding assay used to evaluate ARF binding; the dsDNA is labelled with Cy3 and Atto647N on the opposite sides of the response element (RE). Upon protein binding, the increased distance between the dyes leads to a decrease in FRET efficiency. (b) Titrations of the dsDNA with several ARF variants. The dsDNA alone has a FRET efficiency *E*\* = 0.59; as the protein concentration increases the population of bound DNA (centered at *E*\* = 0.42) increases until all the DNA is bound (saturating condition). A washing step suffices to reset the system proving that the bound population is generated by specific and reversible binding of ARF. Vertical dashed lines are added for visual guidance. (c) Schematic representation of the four-state cyclic model for ARF2-FL KpO-IR7. Note that the dsDNA containing the DNA response element can be found in three states: free (F), bound to a monomer (M) and bound to a dimer (D). The two ARF2 full length variants (K2S and OPCA) have the same DBD (in red) and MR domains (in black) but their PB1 domain (in orange and green, respectively) carry a mutation on either one of the two different surfaces; this hinders oligomerization but allows dimerization. The binding of a dimer to a bipartite response element can occur either through two successive binding events of a monomer or through direct binding of a dimer formed in solution. The dissociation can occur either by the loss of a monomer followed by the dissociation of the second monomer from the DNA or by direct dissociation of a dimer from the DNA.

To demonstrate that the shift seen during the titration is generated by specific binding of ARF to the DNA and that the binding is reversible, we performed a washing step at the end of each titration that reverted the FRET efficiency distributions to the ones seen in absence of ARF. When comparing AtARF2-DBD and AtARF2-FL, the FRET efficiency distributions clearly show the effect of the PB1 domain on the interaction between ARF2 and the IR7 response element; the shift between the response element being mostly free to mostly bound occurs at a protein concentration almost one order of magnitude lower with the full-length protein (256-512 nM ARF2-DBD vs 32-64 nM ARF2-FL). This finding is consistent with the PB1 domain promoting DNA binding. However, the full-length ARF protein also contains an extended MR region. To address the role of the PB1 domain specifically, we engineered mutations in the PB1 domain that prevent head-to tail interaction: AtARF2-FL K2S (K737S) and AtARF2-FL OPCA (D797-8S) carry mutations of amino acids on the positive (K2S) and negative (OPCA) side of the PB1 domain respectively, both of which were shown to impair the interaction between PB1 domains(6). In both mutant ARF2 versions, we see equal percentages of DNA bound and free at concentrations close to the ones of ARF2-DBD (64-128 nM K2S and 128-256 nM OPCA). Thus, PB1 domain interactions contribute to efficient DNA binding. PB1 domains could potentially oligomerize through head-to-tail interactions. To address if oligomerization stabilizes DNA binding, we compared a mixture of ARF2-FL K2S and ARF2-FL-OPCA in a 1:1 ratio (henceforth ARF2 FL KpO) with the wildtype version (ARF2-FL). While the latter allows for oligomerization, the former should only be able to dimerize. Both ARF2 FL KpO and ARF2-FL show high affinity towards the IR7 (32-64 nM). Taken together, these observations indicate that the PB1 domain stabilizes the binding of ARF2 towards an IR7 response element through stabilization of the protein dimer.

### A four-state cyclic model for describing ARF-DNA interactions

One way of quantifying the effect of the PB1 domain on the affinity between ARF and its RE is to fit the increase of the fraction of DNA bound to ARF as the concentration of protein increases, with a single apparent 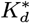. This approach can reliably summarize the strength of the interaction providing a single numerical value that exemplifies at which endogenous protein concentration the interaction becomes relevant(22, 37), but fails to properly describe the underlying system, which results in a lack of predictive power.

The interaction between a protein that can dimerize and a bipartite response element on the DNA can be described using a four-state cyclic model. This model allows for monomers or dimers to bind the DNA, for monomers and dimers to exist in solution and for dimers to form or dissociate both in solution or on the DNA. Figure 1c depicts the model for ARF2-FL KpO; the two ARF2-FL variants are characterized by the same DBD (red) and MR (black) but are mutated on the two opposite surfaces of the PB1 domain (K2S and OPCA mutants in orange and green respectively); this allows for the formation of PB1 domain dimers but hinders the formation of oligomers. The system is defined by four *k*_on_s and four *k*_off_s or alternatively by four equilibrium constants (*K*s). The presence of the PB1 domain should not change the contacting interface between the DBDs and the DNA; hence, the *k*_off_ of the dimer from the DNA (*k*_off,DF_) should have the same value for all AtARF2-variants; the same holds for the *k*_off_ of the monomer from the DNA (*k*_off,MF_). Moreover, the PB1 domain has limited influence on the *k*_on_s of the system which stay diffusion limited. The only constants that are expected to be influenced by changes in the stability of the dimer induced by the PB1 domain are the ones associated with the separation of two monomers: the equilibrium dissociation constant of the dimer in solution (*K*_I_ = *k*_off,DS_*/k*_on,DS_) and *k*_off,DM_, which encompasses the stability of the dimer on the DNA. As the model is a closed cycle, microscopic reversibility(38) implies that only one of these two parameters is a free parameter; then, the different variants tested are characterized in this model by the single variant-specific parameter *K*_I_, which encompasses the stability of the ARF dimer. Fitting experimental results to determine *K*_I_ and the other relevant shared kinetic constants provides a deeper understanding of the system allowing to make prediction for other ARF-AuxRE interactions.

### ARF-IR7 interaction follows a four-state cyclic interaction mechanism

To obtain the kinetics of AtARF2/IR7 interaction, we analyzed relevant datapoints of the titrations for the series of variants using ebFRET(31), a MATLAB suite for Hidden Markov Model (HMM) analysis of single-molecule time traces (see Materials and Methods). As the DNA oligonucleotides are immobilized, their interaction with the proteins in solution can be monitored for several minutes (250 s in our experiments). The analyzed FRET traces returned the most probable hidden state sequence for each trace (Fig. 2a) according to the set of kinetic parameters that best explains the transitions and states seen in the entire dataset.

**Fig. 2.**
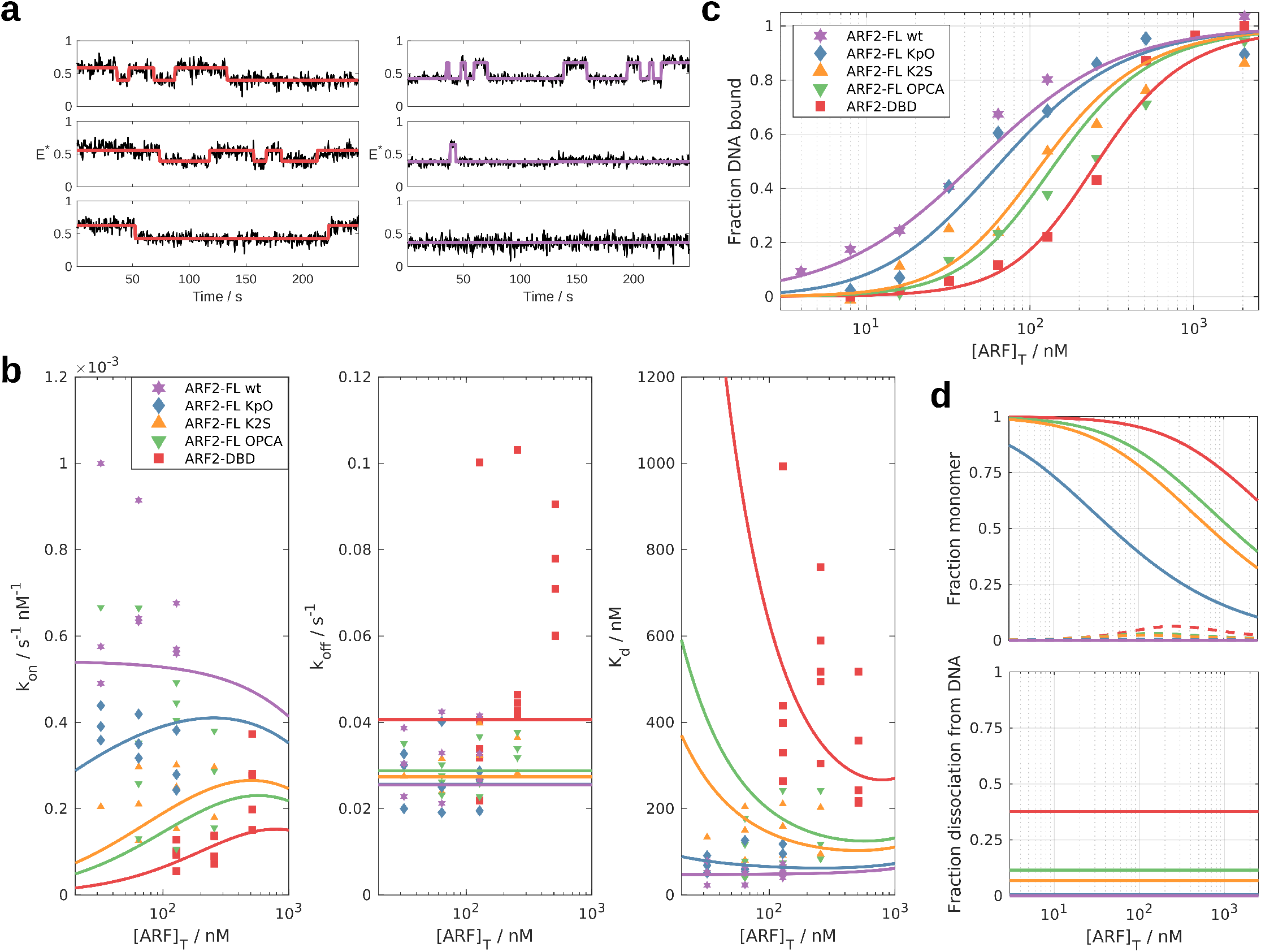
AtARF2-IR7 interaction follows a four-state cyclic interaction mechanism. (a) Example of FRET efficiency time traces of single doubly labelled dsDNA in presence of 128 nM of either AtARF2-DBD (left) or AtARF2-FL wt (right). The FRET efficiency is reported in black and the most probable sequence of hidden states returned by ebFRET is represented in red (AtARF2-DBD) and purple (AtARF2-FL wt). (b) Kinetics parameters obtained from ebFRET. For each ARF variant three concentrations closest to having half of the DNA bound to ARF were measured in at least three independent titrations. Each repeat of each concentration is analyzed independently using ebFRET obtaining a value of observed *k*_on_ and *k*_off_ (and, from their ratio the *K*_d_) and plotted using colored markers. (c) Fraction of DNA bound by ARF as function of ARF concentration (binding isotherm). The fractions bound were obtained from the histograms and plotted using colored markers (Fig. 1b, Materials and Methods). (b-c) The result of the global fit of the kinetics of binding and the fraction of DNA bound is reported as colored lines. (d) Features of the four states system as solved by the global fit. Top: In solution, the monomer is the most abundant species (solid lines). On the DNA, the monomer accounts for less than 10 % of the bound DNA (dashed lines). Bottom: Fraction dissociation of the AtARF2 dimer from the DNA via loss of an AtARF2 monomer. The dissociation of the AtARF2 dimer from the DNA can occur either via its direct unbinding from the DNA or by initial loss of an AtARF2 monomer. Direct unbinding of the dimer is the predominant route for all AtARF2 variants but the fraction of dissociations happening via an initial loss of a monomer accounts for almost 40 % of the events in case of AtARF2-DBD.

To facilitate the identification of transitions by ebFRET, we selected three concentrations of ARF ([ARF]_T_) that returned close to equal populations of ARF-bound and free DNA for each AtARF2 variant. Each AtARF2 variant was tested in at least three independent titrations; each datapoint of each titration was analyzed independently using ebFRET and returned a value for *k*_on_, a value for *k*_off_ and, from their ratio, a value for *K*_d_ (Fig. 2b, colored markers). The observed *k*_on_ show a trend in which ARF variants with higher affinity show faster association. On the other hand, the *k*_off_ show similar values for all the FL variants whilst AtARF2-DBD has a faster *k*_off_. The resulting *K*_d_s show the expected trend, with AtARF2-DBD having the lowest affinity, AtARF2-FL KpO and wt showing the tightest binding and AtARF2-FL K2S and OPCA having an affinity in between these. The trends seen in the *k*_on_ and *k*_off_ suggest that the analysis of the kinetics using HMM is capturing the interaction between the ARF dimer and the DNA and that the interaction between the ARF monomer and the DNA occurs on a timescale shorter than the 500 ms acquisition time used in our experiments.

For a four-states cyclic model the binding isotherms (Fig. 2c, colored markers) and observed kinetic constants (Fig. 2b, colored markers) can be fitted with a system of equations containing a set of three parameters shared across all AtARF2 variants (*k*_on,mic_, *k*_off,DF_ and *k*_off,MF_) and the variant-specific parameter *K*_I_ (see supporting note 1 and 2). Here, *k*_on,mic_ is the microscopic *k*_on_ that a monomer displays when binding a single AuxRE and hence it is equal to *k*_on,MM_ but it is half the value of *k*_on,M_ and *k*_on,D_. The global fit (Fig. 2b-c, colored lines) returned the values of *k*_on,mic_, *k*_off,DF_, *k*_off,MF_ and *K*_I_s that best explain the experimental data (see Table 1). The global fit converged to a *K*_I_ of 0 nM for AtARF2-FL wt; in this situation the equation of the fraction bound for the four-states system simplifies to a simple binding isotherm for the dimer (see supporting note 1). On the other binding isotherms, the fit captured the shift of the binding to higher [ARF]_T_ thanks to increasing values of *K*_I_, which corresponds to a decrease in dimer stability (Fig. 2c). The fit of the binding isotherm of AtARF2FL KpO is still very close to the one of AtARF2FL wt but because of the decrease in dimer stability (*K*_I_ = 0.016 µM) its steepness is increased. The two AtARF2-FL mutants, K2S and OPCA, show similar values of *K*_I_ (0.23 µM and 0.41 µM respectively). Lastly, the fit returned a value of *K*_I_ of 1.9 µM for AtARF2-DBD.

**Table 1.** Global fit: values and uncertainty of the fitting parameters reported as mean [95 % CI].

| Protein | $k_{on,mic}$<br>[nM<sup>-1</sup>s<sup>-1</sup>] | $k_{off,MF}$<br>[s<sup>-1</sup>] | $k_{off,DF}$<br>[s<sup>-1</sup>] | $K_d$<br>[μM] |
| --- | --- | --- | --- | --- |
| AtARF2-DBD |  |  |  | 1.9 [0.9:2.9] × 10<sup>0</sup> |
| AtARF2-FL OPCA |  |  |  | 4.1 [1.0:7.1] × 10<sup>-1</sup> |
| AtARF2-FL K2S | 5.4 [3.6:7.3] × 10<sup>-4</sup> | 1.7 [0.6:2.8] | 2.6 [2.2:2.9] × 10<sup>-2</sup> | 2.3 [0.4:4.2] × 10<sup>-1</sup> |
| AtARF2-FL KpO |  |  |  | 1.6 [-1.5:4.7] × 10<sup>-2</sup> |
| AtARF2-FL wt |  |  |  | 0.0 [-0.4:0.4] × 10<sup>-6</sup> |

Looking at the observed binding kinetics, the global fit captures the trends of the observed *k*_on_ and *k*_off_ (Fig. 2b). Here, AtARF2 variants with higher dimer stability display higher values of observed *k*_on_ as their lower *K*_I_ increases the effective concentration of ARF dimer in solution. The fits for the observed *k*_off_ of AtARF2-FL wt and KpO converge to the value of the dissociation kinetic of the dimer from the DNA (*k*_off,DF_ = 0.026 s^−1^). For the other datasets (AtARF2-FL K2S, AtARF2-FL OPCA and AtARF2-DBD) the dissociation of the dimer from the DNA caused by the loss of a monomer plays a role and becomes almost as likely as the dissociation of the dimer from the DNA in the case of AtARF2-DBD (*k*_off,DM_ = 0.016 s^−1^).

The kinetic and equilibrium constants obtained from the global fit show that the monomer is the predominant species in solution for most of AtARF2 variants and for most of the tested concentration range (Fig. 2d top, solid lines). Strikingly, the fraction of DNA bound by a monomer never exceeds 10 % (Fig. 2d top, dashed lines). The complex consisting of the AtARF2 dimer bound to the DNA can split by either a monomer or the dimer dissociating from the DNA. Our results show that the dissociation of the dimer from the DNA is the route used by AtARF2-FL and AtARF2-FL KpO, while the dissociation from the DNA through loss of a monomer becomes viable for AtARF2-FL K2S and OPCA and accounts for approximatively 40 % of the splitting events in the case AtARF2-DBD (Fig. 2d bottom).

### Dimer stability determines the binding kinetics of ARF-DBDs

We showed that AtARF2 dimer stability induced by the presence of the PB1 domain influences the kinetics of the binding of ARF towards its DNA response element. We next asked whether dimer stability is a more generic parameter defining DNA binding affinity across the ARF protein family. To this end, we purified the DNA-binding domains of two other ARFs (class B AtARF1-DBD, class A AtARF5-DBD) and two mutant versions (AtARF5-DBD G279N and AtARF5-DBD R215A) and quantified their DNA binding affinity.

Experiments with AtARF1-DBD showed similar values of *k*_on_ and *k*_off_ (1.3 × 10^−4^ nM^−1^s^−1^ 95 % CI [0.8:1.8], 0.080 s^−1^ 95 % CI [0.062:0.098], respectively) as AtARF2-DBD (1.1 × 10^−4^ nM^−1^s^−1^ 95 % CI [0.7:1.4], 0.066 s^−1^ 95 % CI [0.045:0.087], respectively; Fig. 3). On the other hand, AtARF5-DBD showed a 5-fold increase in *k*_on_ (5.9 × 10^−4^ nM^−1^s^−1^ 95 % CI [2.3:9.4]) and an 8-fold reduction in *k*_off_ (0.0085 s^−1^ 95 % CI [0.0051:0.0118]) compared to AtARF2-DBD; which lead to a *K*_d_ of 15 nM (95 % CI [12:18]). In analogy with the considerations made for AtARF2-FL, the increase in *k*_on_ and part of the decrease in *k*_off_ can be explained with AtARF5-DBDs forming a tighter protein dimer compared to AtARF1 and AtARF2 DBDs.

**Fig. 3.**
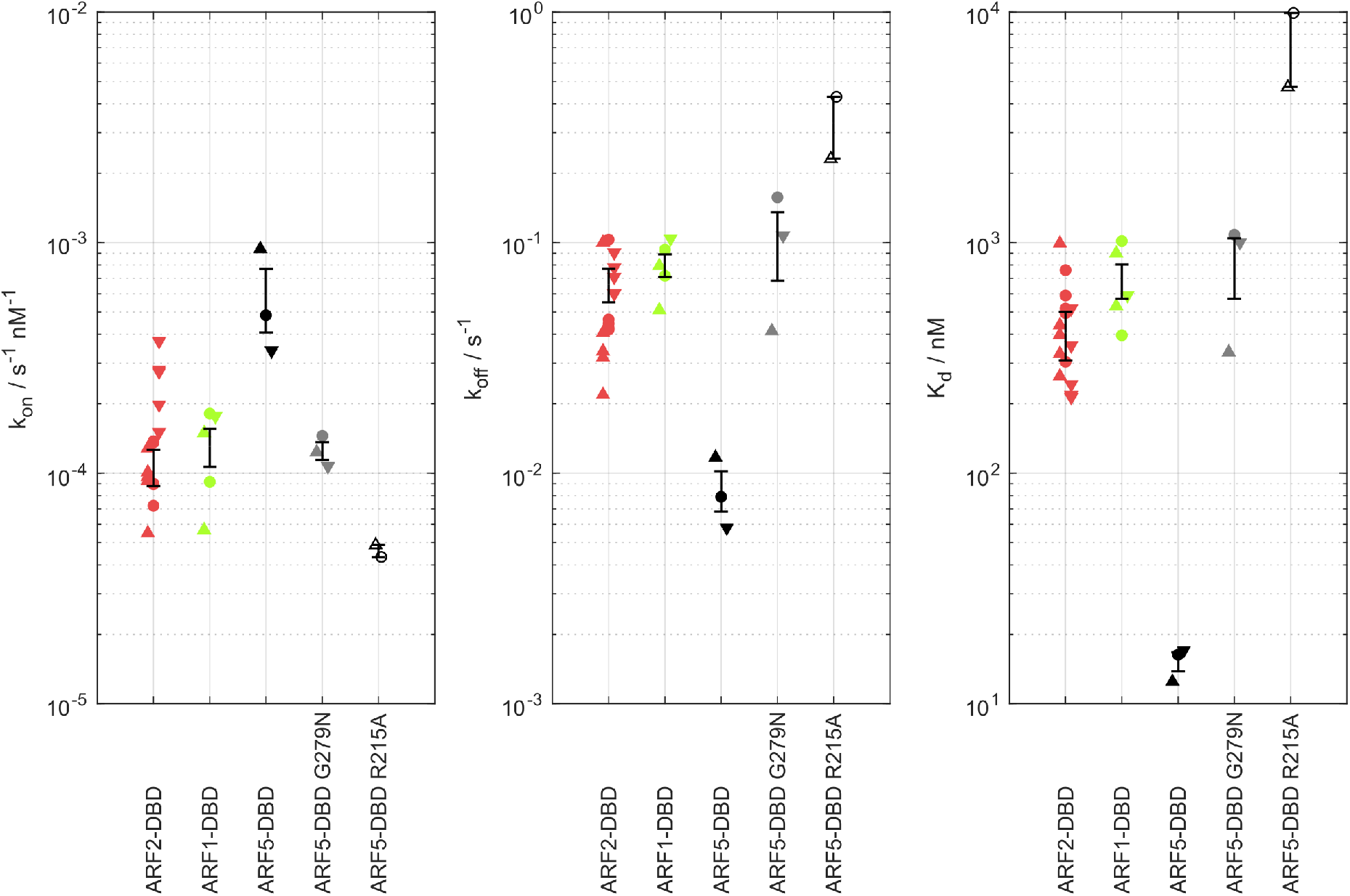
Kinetics of the interaction between AtARF-DBDs and IR7. Kinetic parameters obtained from HMM analysis using ebFRET. The datapoints are marked with Δ, □ and ∇ in order of increasing ARF concentration. The ARF concentrations were 128, 256, 512 nM for AtARF2-DBD and AtARF1-DBD, 8, 16 and 32 nM for AtARF5-DBD, 64, 128 and 256 nM for AtARF5-DBD G279N and 128 and 512 nM for AtARF5-DBD R215A. The error bars represent the standard deviations of the mean values. AtARF2-DBD and AtARF1-DBD behaved similarly while AtARF5-DBD showed increased *k*_on_ and decreased *k*_off_. Consistent with a model in which part of the difference in kinetic can be explained by an increased stability of AtARF5-DBD dimer. A weakening of AtARF5-DBD dimerisation (G279N mutant) leads to kinetic parameters that resemble the ones of AtARF1-DBD and AtARF2-DBD. In addition, AtARF5-DBD R215A mutant in a key amino acid for the interaction with DNA showed a *k*_on_ reduced by one order of magnitude and a *k*_off_ increased by almost two orders of magnitude compared to the wild-type.

To test this hypothesis directly, we tested AtARF5-DBD G279N, a single amino acid mutation known to reduce AtARF5-DBD dimerization(17). Strikingly, the kinetics of the interaction between AtARF5-DBD G279N and the IR7 became similar to the ones of AtARF1 and AtARF2 DBDs (1.2 × 10^−4^ nM^−1^s^−1^ 95 % CI [1.0:1.5], 0.10 s^−1^ 95 % CI [0.04:0.17]) validating our hypothesis. Finally, we tested AtARF5-DBD R215A, a mutant in which a key amino acid for the interaction with the DNA is mutated(17). This mutant showed a 13-fold reduction of *k*_on_ compared to the wild-type (0.46 × 10^−4^ nM^−1^s^−1^ 95 % CI [0.41:0.51] as well as a 39-fold increase of *k*_off_ (0.33 s^−1^ 95 % CI [0.14:0.52]) which translates in a reduction of affinity of three orders of magnitude. We note that the magnitude of the reduction of *k*_on_ is consistent with the effect of charge neutralization of DNA-contacting residues seen in other protein-DNA interactions(39) and is a reminder of the importance of charged residues in defining association kinetics (40, 41).

To directly measure ARF dimer stability, we measured SAXS (Small-angle X-ray scattering) intensity profiles of AtARF5-DBD and AtARF1-DBD (Fig. 4). The difference in dimer stability is clear with AtARF5-DBD exhibiting higher dimer prevalence at all tested concentration. The results on AtARF5-DBD are consistent with a dimerization *K*_d_ in the order of the tenth of µM while for AtARF1-DBD the *K*_d_ is in the order of few µM. These results confirm the expectation set by the analysis of the binding kinetics that AtARF5-DBD forms relatively stable dimers even in absence of the PB1 domain.

**Fig. 4.**
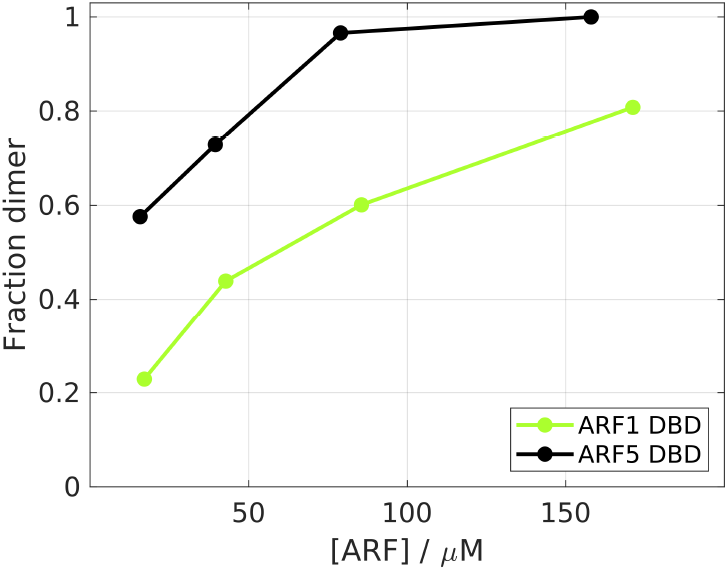
Fraction of dimer measured using SAXS. The stability of the dimer of AtARF5-DBD is higher than the one of AtARF1-DBD for all tested concentration.

## Discussion

The ARF PB1 domain mediates the binding of Aux/IAA to ARFs, allowing for inhibition of ARF activity(3–5). In ARFs from Arabidopsis, deleting or mutating the PB1 domain leads to hyperactive ARFs(23), consistent with a role in suppressing activity. In contrast, the Marchantia ARF1 PB1 domain is required for function, a deletion of this domain renders the protein inactive(22). Given that the minimal set of ARF proteins found in Marchantia qualifies these as likely representatives of ancestral protein functions(42), an open question is what the actual roles of ARF PB1 domains are. Here, we explored the role of this domain in modulating the DNA binding affinity of AtARF2 towards an IR7 response element. We found that full-length ARF2 protein has a strongly increased DNA-binding affinity, which can be ascribed to interactions between the PB1 domains. Interestingly, our results show that oligomerization does not further enhance the affinity towards bipartite response elements. This behaviour is consistent with the fact that additional ARF monomers (beside the initial two) do not have any AuxRE left to further stabilize the binding to the DNA. This said, the effect of the PB1 domain seen in our experiments predicts that oligomerization should be relevant on response elements comprising of more than two AuxREs as the PB1 domain would enable cooperative binding beyond the dimer. Given the short consensus sequence in AuxRE motifs, these may occur in close proximity in promoters, in which case oligomerization could generate additional cooperativity of ARF-DNA interaction. Regardless, the use of a C-terminal head-to tail oligomerization domain (with two interaction faces) can be considered an efficient means of flexible interaction.

By simultaneously fitting affinity and kinetic data with the analytical solution of a four-state cyclic model, we showed that the increase of ARF DNA binding affinity can be completely attributed to a shift in the dimer/monomer equilibrium of ARFs. It follows that the effect of the PB1 domain on ARFs affinity towards an IR7 response element is to shift the dimerization equilibrium towards the dimer. Strikingly, the fit allows to obtain quantitative information about the protein dimerization *K*_d_ in solution (i.e., *K*_I_) although the protein was not directly observed in the experiment.

The kinetic parameters for AtARF2-DBD show that DNA binding and unbinding is almost equally probable to happen through a monomer-bound DNA intermediate or through direct binding of the dimer (Fig. 5). Moreover, the monomer is the most common species in solution even at ARF concentrations that saturate the DNA (i.e., DNA fully bound by an ARF dimer); despite this, the percentage of DNA bound by an ARF monomer never exceeds 10 % as this intermediate is short lived and quickly proceeds to either forming a dimer or dissociate.

**Fig. 5.**
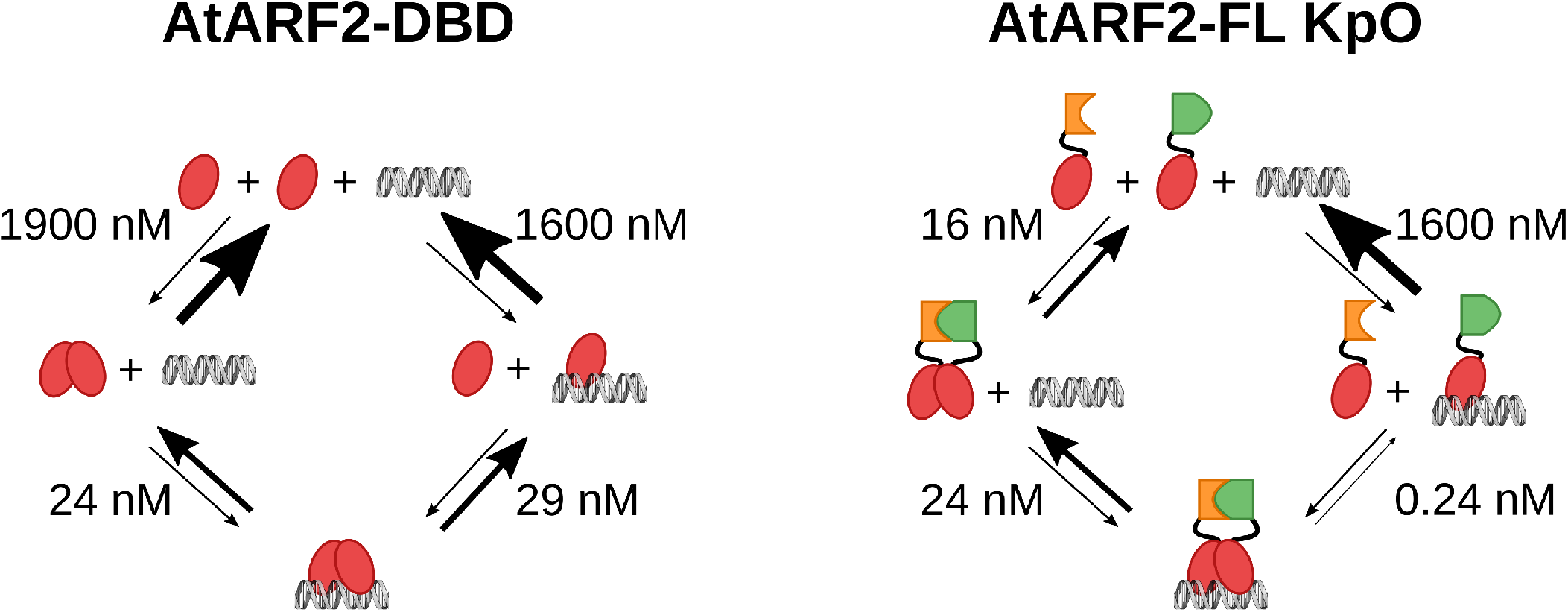
Dissociation constants for the four-state cyclic model as determined from the fit in figure 2. ARF2-DBD binds the DNA via dimerization on the DNA and dimerization in solution with similar probability. ARF2-FL KpO mostly dimerizes in solution and then binds the DNA. The presence of the PB1 domain reduces all Kds making all interactions more stable (aside from the monomer-DNA interaction). This results in the higher affinity of ARF towards the RE when the PB1 domain is present.

The kinetics of AtARF2-FL KpO-IR7 interaction is remarkably different (Fig. 5); here, the association between the DNA and a dimer follows almost exclusively the pathway where the ARF dimer is formed in solution. Moreover, the unbinding of a single monomer from a dimer bound to the DNA is unlikely (*K*_d_ *<* 1 nM).

In general, the importance of dimerization for stable DNA binding clearly emerges from our analysis as a reminder of the importance of cooperativity in protein-DNA interaction. Moreover, cooperativity is symmetric: a protein that can dimerize on a bipartite response element will bind it with higher affinity but also a bipartite element will stabilize the dimer of the protein that is bound to it. In particular, the dimer of AtARF2 is ≈ 60 times more stable when bound to the DNA compared to being in solution.

The analysis of the kinetics of the interaction between different ARF DBDs and the IR7-RE suggests that the tighter binding of AtARF5-DBD compared to AtARF1 and AtARF2 DBDs is in part due to the higher stability of its dimer. This prediction is further corroborated by SAXS data showing that AtARF5-DBD forms more stable dimers in solution compared to AtARF1-DBD. The stable DNA binding that AtARF5-DBD achieve even in absence of the PB1 domain could explain why AtARF5PB1 is a gain-of-function mutant that can activate auxin-responsive genes even in absence of auxin(23). Then, the role of the PB1 domain of AtARF5 appears to be mainly to bind the PB1 domain of Aux/IAAs coupling the transcriptional output of ARF with the presence of auxin. This hypothesis is confirmed by the fact that the PB1 domain of AtARF5 has a homodimerization *K*_d_ of 870 nM but an heterodimerization *K*_d_ with the PB1 domain of Aux/IAA17 of 73 nM(8). Moreover, AtARF5 and other A-class ARFs have been found to interact with many different Aux/IAA in a series of protein-protein interaction assays(24, 43–53). A different scenario is seen in case of AtARF2 (a class B ARF), where our data suggest that the interaction between PB1 of different AtARF2 monomers might be required to achieve the stable DNA binding that enables protein function. This behavior of the PB1 domain might be a common feature of other class B/C ARFs and could explain why this class of ARFs has been seen to interact with fewer members of the Aux/IAA family(43, 51–54).

The picture emerging is that the PB1 domain has different functions in the two main ARF classes in *A. thaliana*. In class A ARFs, it serves as a mediator of auxin-responsiveness whereas it stabilizes the DNA binding in class B (and perhaps C) ARFs. This model of action for the PB1 domain is similar to the one found in *M. polymorpha* as part of the recently published minimal auxin response system(22) with one key difference: MpARF1 (the only class A ARF in this species) cannot function without its PB1 domain. Therefore, the PB1 domain of class A ARFs in *M. polymorpha* probably has the double function of stabilizing the binding to the DNA and interacting with MpAux/IAA. This double function opens the possibility of a double repression by Aux/IAA, where, in addition to the recruitment of the co-repressor TOPLESS(55) (TPL), a destabilization of ARF-DNA interaction might also play a role.

Binding of ARF to bipartite AuxREs in the other two possible orientations (directed repeat DR and everted repeat ER), should resemble the one seen for the IR with the difference that the dimerization through the DBD domain should not be possible. In this scenario the analytical solution of the 4-state model for ARF-DBDs simplifies to a binding isotherm for independent binding of the monomers characterized by low steepness (no cooperativity, see also supporting Note 1). Strikingly, titrations presented in a recent publication(37) confirmed this prediction; the binding of AtARF1-DBD and AtARF5-DBD to a bipartite DR5 was compatible with a simple binding isotherm, whereas binding to an IR8 showed steeper response, similar to the one seen here for AtARF2-DBD. Since stable DNA binding arises from stable dimerization/oligomerization, the topology of composite AuxREs dictates the affinity towards distinct ARF members differentiating them based on the relative strength of the homotypic interaction through their DBD and PB1 domains. Tweaking the affinities of different ARFs towards the same DNA sequence can be achieved by affecting their dimerization properties and opens the possibility for an evolutionary pathway of complex interactions between members of the family.

Lastly, it is interesting to speculate on the biological significance of the dual, cooperative dimerization mode we identified here. Effectively, the double-check mechanism would favor dimerization of ARFs, when bound on DNA. ARF monomers have limited sequence specificity of DNA binding. In the hexanucleotide binding site, only 2 nucleotides are invariant, and four are conserved(17, 19, 56). Thus, one would expect a monomer to find binding sites frequently in the genome. Dimerization adds two constraints that dramatically increase specificity: a second, symmetric DNA element as well as a fixed optimal space between the elements. This strongly limits the probability of a random occurrence of the response element and explains why dimerization is such a common features of transcription factors across all domains of life(57, 58). Unfortunately, probing genome-wide ARF binding has been challenging, and no comparisons between monomers and dimers have yet been made. However, ARF2 and ARF5 have both been used in DAP-seq binding site mapping on the Arabidopsis genome(19). If dimerization limits the number of genomic binding sites, one would predict that ARF5 – with a higher propensity to dimerize (as shown here) binds fewer sites. This is exactly what was found: ARF2 appears more promiscuous in its binding profile(19, 56). A hypothesis, to be tested in the future, is therefore that dimerization is the primary mechanism for defining ARF-DNA binding specificity in vivo.

## Authors’ Contributions

The project was initiated and supervised by Dolf Weijers (D.W.) and Johannes Hohlbein (J.H.). Mattia Fontana (M.F.) developed the methodology, defined the experimental design, wrote and adapted the software for data analysis. Mark Roosjen and Willy van den Berg purified the proteins. M.F. performed smFRET experiments, analyzed the data, derived the analytical solutions for the system and implemented the global fit. Isidro Crespo García performed SAXS experiments and data analysis, supported by Marc Malfois and under supervision by Roeland Boer. M.F., J.H. and D.W discussed the content of the manuscript. M.F. wrote the draft manuscript and produced the figures. All authors provided feedback on the draft.

## Data availability

The experimental data is available on Zenodo: https://doi.org/10.5281/zenodo.7249508.

## Acknowledgements

We would like to thank all our colleagues at the Laboratory of Biophysics and the Laboratory of Biochemistry for helpful discussions.

