## Supporting Information for "Cooperative action of separate interaction domains promotes high-affinity DNA binding of *Arabidopsis thaliana* ARF transcription factors"

DRAFT

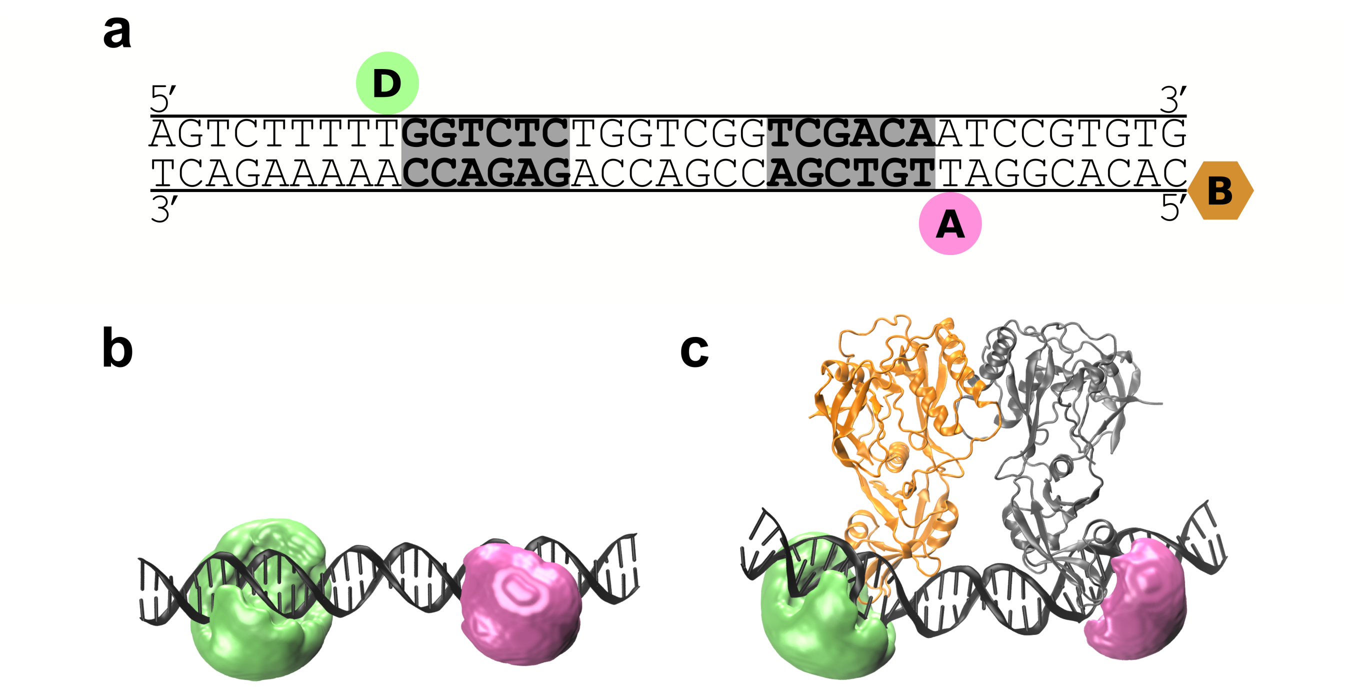


**Fig. S1.** dsDNA oligo and accessible volumes (AVs). (a) Schematic representation of the dsDNA oligo; donor (Cy3, green) and acceptor (ATTO647N, magenta) indicate the positions for internal labelling; similarly, the biotin (brown) used to immobilize the dsDNA is conjugated to a modified 5'- end. The binding sites for ARF (AuxREs) are highlighted in gray. (b-c) Accessible volumes (AVs) for Cy3 and ATTO647N in absence (b) and presence (c) of AtARF1-DBD. The AVs show the volume that a fluorophore can explore thanks to its flexible linker. The parts of the AV facing each other become partially inaccessible to the dyes upon protein binding; this effect accounts for ($\approx 56\,\%$) of the expected distance change between the mean position of the dyes, with DNA bending having a smaller effect ($\approx 44\,\%$ of the total change). The fluorophores being on average further apart when the protein is present leads to a decrease in FRET efficiency.

DRAFT

### Supporting Note 1: Derivation binding isotherm four states system

The dsDNA containing the DNA response element can be found in three states: free (F), bound to a monomer (M) and bound to a dimer (D). The DNA oligo has a certain probability $P$ to be in each state and can transition to a different state by either binding or dissociating from a monomer or a dimer. Applying the law of mass action allows to write the following three ordinary differential equations, which can further be simplified assuming the system is in equilibrium

$$\cancelto{0}{\frac{dP_{\mathrm{F}}}{dt}} = k_{\mathrm{off,DF}} * P_{\mathrm{D}} + k_{\mathrm{off,MF}} * P_{\mathrm{M}} - k_{\mathrm{on,M}} * [\mathrm{ARF}] * P_{\mathrm{F}} - k_{\mathrm{on,D}} * [\mathrm{ARF}_2] * P_{\mathrm{F}} \quad [1]$$

$$\cancelto{0}{\frac{dP_{\mathrm{M}}}{dt}} = k_{\mathrm{on,M}} * [\mathrm{ARF}] * P_{\mathrm{F}} + k_{\mathrm{off,DM}} * P_{\mathrm{D}} - k_{\mathrm{off,MF}} * P_{\mathrm{M}} - k_{\mathrm{on,MM}} * [\mathrm{ARF}] * P_{\mathrm{M}} \quad [2]$$

$$\cancelto{0}{\frac{dP_{\mathrm{D}}}{dt}} = k_{\mathrm{on,D}} * [\mathrm{ARF}_2] * P_{\mathrm{F}} + k_{\mathrm{on,MM}} * [\mathrm{ARF}] * P_{\mathrm{M}} - k_{\mathrm{off,DM}} * P_{\mathrm{D}} - k_{\mathrm{off,DF}} * P_{\mathrm{D}}. \quad [3]$$

Moreover, the sum of the probabilities of the DNA being in any of the three states must be unity

$$P_{\mathrm{F}} + P_{\mathrm{M}} + P_{\mathrm{D}} = 1. \quad [4]$$

Assuming that the fraction of ARF bound to the DNA is negligible compared to the one of ARF in solution, it is possible to derive the following equation linking the concentration of dimers in solution with the total concentration of ARF added to the solution

$$[\mathrm{ARF}_2] = \frac{4[\mathrm{ARF}]_{\mathrm{T}} + K_{\mathrm{I}} - \sqrt{K_{\mathrm{I}} * (8[\mathrm{ARF}]_{\mathrm{T}} + K_{\mathrm{I}})}}{8} \quad [5]$$

where $K_{\mathrm{I}}$ is the dissociation constant of the protein dimer (in solution) into two monomers. Since the four states system forms a closed circle, one of the kinetic constant must be obtained from the equation imposing microscopic reversibility(1)

$$k_{\mathrm{off,DM}} = K_{\mathrm{I}} \frac{k_{\mathrm{on,M}} * k_{\mathrm{on,MM}} * k_{\mathrm{off,DF}}}{k_{\mathrm{off,MF}} * k_{\mathrm{on,D}}}. \quad [6]$$

The equations 1 to 6 form a system that can be simplified by substitution to obtain the equations for the expected $P_{\mathrm{M}}$ and $P_{\mathrm{D}}$ as function of $[\mathrm{ARF}]_{\mathrm{T}}$. Moreover, we can define the system as being function of a single microscopic association kinetic constant $k_{\mathrm{on,mic}}$, which represents the kinetics of the association of a monomer to a single AuxRE. Then, the three $k_{\mathrm{on}}$s of the four states model can be defined in relation to this single constant ($k_{\mathrm{on,MM}} = k_{\mathrm{on,mic}}$, $k_{\mathrm{on,M}} = 2k_{\mathrm{on,mic}}$ and $k_{\mathrm{on,D}} = 2k_{\mathrm{on,mic}}$).

$$P_{\mathrm{M}} = \frac{8k_{\mathrm{on,mic}}k_{\mathrm{off,DF}}K_{\mathrm{I}}\left(\left(K_{\mathrm{I}}k_{\mathrm{on,mic}}+k_{\mathrm{off,MF}}\right)A-\frac{k_{\mathrm{on,mic}}B}{2}+\left(-\frac{K_{\mathrm{I}}}{2}+2[\mathrm{ARF}]_{\mathrm{T}}\right)k_{\mathrm{on,mic}}-k_{\mathrm{off,MF}}\right)}{\begin{array}{r} 8k_{\mathrm{on,mic}}\left(-\frac{k_{\mathrm{off,MF}}k_{\mathrm{on,mic}}B}{8}+\left(\left(k_{\mathrm{off,DF}}-\frac{3k_{\mathrm{off,MF}}}{8}\right)K_{\mathrm{I}}+\frac{[\mathrm{ARF}]_{\mathrm{T}}k_{\mathrm{off,MF}}}{2}\right)k_{\mathrm{on,mic}}+\frac{3k_{\mathrm{off,DF}}k_{\mathrm{off,MF}}}{2}\right)K_{\mathrm{I}}A- \\ -4K_{\mathrm{I}}\left(\left(k_{\mathrm{off,DF}}-\frac{3k_{\mathrm{off,MF}}}{4}\right)K_{\mathrm{I}}-4[\mathrm{ARF}]_{\mathrm{T}}\left(k_{\mathrm{off,DF}}+\frac{3k_{\mathrm{off,MF}}}{4}\right)\right)k_{\mathrm{on,mic}}^2+16k_{\mathrm{off,DF}}k_{\mathrm{off,MF}}^2- \\ -4k_{\mathrm{on,mic}}\left(K_{\mathrm{I}}\left(k_{\mathrm{off,DF}}-\frac{k_{\mathrm{off,MF}}}{4}\right)k_{\mathrm{on,mic}}+k_{\mathrm{off,MF}}^2\right)B+ \\ +4\left(\left(k_{\mathrm{off,DF}}+k_{\mathrm{off,MF}}\right)K_{\mathrm{I}}+4[\mathrm{ARF}]_{\mathrm{T}}k_{\mathrm{off,MF}}\right)k_{\mathrm{off,MF}}k_{\mathrm{on,mic}} \end{array}}, \quad [7]$$

$$P_{\mathrm{D}} = \frac{-3k_{\mathrm{on,mic}}\left(K_{\mathrm{I}}k_{\mathrm{on,mic}}\left(K_{\mathrm{I}}-\frac{4[\mathrm{ARF}]_{\mathrm{T}}}{3}+\frac{B}{3}\right)A+\left(\frac{4k_{\mathrm{off,MF}}}{3}-\frac{K_{\mathrm{I}}k_{\mathrm{on,mic}}}{3}\right)B-(K_{\mathrm{I}}+4[\mathrm{ARF}]_{\mathrm{T}})\left(K_{\mathrm{I}}k_{\mathrm{on,mic}}+\frac{4k_{\mathrm{off,MF}}}{3}\right)\right)k_{\mathrm{off,MF}}}{\begin{array}{r} 8k_{\mathrm{on,mic}}\left(-\frac{k_{\mathrm{off,MF}}k_{\mathrm{on,mic}}B}{8}+\left(\left(k_{\mathrm{off,DF}}-\frac{3k_{\mathrm{off,MF}}}{8}\right)K_{\mathrm{I}}+\frac{[\mathrm{ARF}]_{\mathrm{T}}k_{\mathrm{off,MF}}}{2}\right)k_{\mathrm{on,mic}}+\frac{3k_{\mathrm{off,DF}}k_{\mathrm{off,MF}}}{2}\right)K_{\mathrm{I}}A- \\ -4K_{\mathrm{I}}\left(\left(k_{\mathrm{off,DF}}-\frac{3k_{\mathrm{off,MF}}}{4}\right)K_{\mathrm{I}}-4[\mathrm{ARF}]_{\mathrm{T}}\left(k_{\mathrm{off,DF}}+\frac{3k_{\mathrm{off,MF}}}{4}\right)\right)k_{\mathrm{on,mic}}^2+16k_{\mathrm{off,DF}}k_{\mathrm{off,MF}}^2- \\ -4k_{\mathrm{on,mic}}\left(K_{\mathrm{I}}\left(k_{\mathrm{off,DF}}-\frac{k_{\mathrm{off,MF}}}{4}\right)k_{\mathrm{on,mic}}+k_{\mathrm{off,MF}}^2\right)B+ \\ +4\left(\left(k_{\mathrm{off,DF}}+k_{\mathrm{off,MF}}\right)K_{\mathrm{I}}+4[\mathrm{ARF}]_{\mathrm{T}}k_{\mathrm{off,MF}}\right)k_{\mathrm{off,MF}}k_{\mathrm{on,mic}} \end{array}}, \quad [8]$$

where $A = \sqrt{(8[\mathrm{ARF}]_{\mathrm{T}} + K_{\mathrm{I}})/K_{\mathrm{I}}}$ and $B = \sqrt{K_{\mathrm{I}}(8[\mathrm{ARF}]_{\mathrm{T}} + K_{\mathrm{I}})}$. Then, $P_{\mathrm{M}}$ and $P_{\mathrm{D}}$ are defined by 4 parameters: $k_{\mathrm{on,mic}}$, $k_{\mathrm{off,MF}}$, $k_{\mathrm{off,DF}}$ and $K_{\mathrm{I}}$. The experimental determination of the fraction bound is based on fitting two Gaussians at the $E^*$s of the free and dimer-bound DNA; the monomer is expected to reside halfway between these two populations and it is reasonable to assume that half of the monomer bound population will be erroneously accounted as free and half as bound. Following this assumption we can define the fraction bound as

$$F_{\mathrm{B}} = P_{\mathrm{D}} + \frac{1}{2}P_{\mathrm{M}}. \quad [9]$$

DRAFT

In the limit of an infinitely stable protein dimer (i.e., $K_\mathrm{I} = 0$), the function for $F_\mathrm{B}$ (9) simplifies to the simple binding isotherm for the direct binding (and unbinding) of a dimer to the DNA

$$F_\mathrm{B} = \frac{[\mathrm{ARF}]_\mathrm{T}}{[\mathrm{ARF}]_\mathrm{T} + 2\frac{k_\mathrm{off,DF}}{k_\mathrm{on,D}}}. \qquad [10]$$

When $K_\mathrm{I} > 0$ the presence of monomers in solution allows for the association (and dissociation) of the dimer to the DNA to occur via a second route; this path passes through an intermediate state where a monomer is bound to the DNA. As the stability of the dimer in solution decreases (i.e., as $K_\mathrm{I}$ increases), this second route for the association (and dissociation) of a protein dimer to the DNA becomes more and more likely. For sufficiently high values of $K_\mathrm{I}$, the association (or dissociation) of a dimer from the DNA happens almost exclusively via successive binding (or unbinding) of two monomers; in this situation, the function for $F_\mathrm{B}$ (9) converges to the Klotz equation(2)

$$F_\mathrm{B,Kl} = \frac{1}{2}\frac{K_1[ARF] + 2K_1K_2[ARF]^2}{1 + K_1[ARF] + K_1K_2[ARF]^2} \qquad [11]$$

where $K_1 = k_\mathrm{on,M}/k_\mathrm{off,MF}$ and $K_2 = k_\mathrm{on,MM}/k_\mathrm{off,DM}$. $K_1$ and $K_2$ are stoichiometric association constants for the first and second binding event respectively. For independent binding (i.e., no cooperativity), the Klotz equation simplifies to a simple binding isotherm in which the dissociation constant is defined by the microscopic kinetic constants related to the interaction of one monomer to one binding site ($K_\mathrm{d,mic} = k_\mathrm{off,mic}/k_\mathrm{on,mic}$).

Summarizing the behaviour of the binding isotherm of the four-state cyclic model (Eq. 9), at values of $K_\mathrm{I}$ close to zero it approximates the simple binding isotherm of the dimer, for intermediate values of $K_\mathrm{I}$ the function is steeper capturing the expected cooperativity, while at high values of $K_\mathrm{I}$ the function approximates the Klotz equation and its steepness ultimately decreases to that of a simple binding isotherm for independent binding.

DRAFT

### Supporting Note 2: Derivation of observed $k_{\mathrm{on}}$ and $k_{\mathrm{off}}$ four states system

Given the four state model we can write the following notable transition probabilities involving changes in the binding state of the DNA

$$
\begin{aligned}
P_{\mathrm{F}\mapsto\mathrm{M}}(t) &= \frac{k_{\mathrm{on,M}}}{FR} * [\mathrm{ARF}] * e^{-\left(\frac{k_{\mathrm{on,D}}}{\mathrm{FR}} * [\mathrm{ARF}_2] + \frac{k_{\mathrm{on,M}}}{FR} * [\mathrm{ARF}]\right)t} \\
P_{\mathrm{M}\mapsto\mathrm{D}}(t) &= \frac{k_{\mathrm{on,MM}}}{\mathrm{FR}} * [\mathrm{ARF}] * e^{-\left(\frac{k_{\mathrm{off,MF}}}{\mathrm{FR}} + \frac{k_{\mathrm{on,MM}}}{\mathrm{FR}} * [\mathrm{ARF}]\right)t} \\
P_{\mathrm{F}\mapsto\mathrm{D}}(t) &= \frac{k_{\mathrm{on,D}}}{\mathrm{FR}} * [\mathrm{ARF}_2] * e^{-\left(\frac{k_{\mathrm{on,D}}}{\mathrm{FR}} * [\mathrm{ARF}_2] + \frac{k_{\mathrm{on,M}}}{\mathrm{FR}} * [\mathrm{ARF}]\right)t} \\
P_{\mathrm{D}\mapsto\mathrm{M}}(t) &= \frac{k_{\mathrm{off,DM}}}{\mathrm{FR}} * e^{-\left(\frac{k_{\mathrm{off,DF}}}{\mathrm{FR}} + \frac{k_{\mathrm{off,DM}}}{\mathrm{FR}}\right)t} \\
P_{\mathrm{D}\mapsto\mathrm{F}}(t) &= \frac{k_{\mathrm{off,DF}}}{\mathrm{FR}} * e^{-\left(\frac{k_{\mathrm{off,DF}}}{\mathrm{FR}} + \frac{k_{\mathrm{off,DM}}}{\mathrm{FR}}\right)t},
\end{aligned}
\qquad [12]
$$

where FR is the frame rate in fps ($\mathrm{fs}^{-1}$) and the time $t$ is expressed in frames (f). Since the monomer is expected to have short dwell time on the DNA (compared to the frame time of 500 ms), the only transitions that can be observed are the ones that lead to a protein dimer bound to the DNA. Then, the observed transition probability associated with "binding" and "dissociation" can be derived as

$$P_{\mathrm{F,M}\mapsto\mathrm{D}} = \frac{P_{\mathrm{F}}}{P_{\mathrm{F}} + P_{\mathrm{M}}}(P_{\mathrm{F}\mapsto\mathrm{D}} + P_{\mathrm{F}\mapsto\mathrm{M}} * P_{\mathrm{M}\mapsto\mathrm{D}} * \tau_{\mathrm{M}}) + \frac{P_{\mathrm{M}}}{P_{\mathrm{F}} + P_{\mathrm{M}}} P_{\mathrm{M}\mapsto\mathrm{D}}, \qquad [13]$$

$$P_{\mathrm{D}\mapsto\mathrm{F,M}} = P_{\mathrm{D}\mapsto\mathrm{F}} + P_{\mathrm{D}\mapsto\mathrm{M}}, \qquad [14]$$

where $\tau_{\mathrm{M}}$ is the dwell time of the monomer on the DNA ($\tau_{\mathrm{M}} = \mathrm{FR}/(k_{\mathrm{off,MF}} + k_{\mathrm{on,MM}} * [\mathrm{ARF}])$). Then, these probabilities can be converted to the observed $k_{\mathrm{on}}$ and $k_{\mathrm{off}}$

$$k_{\mathrm{on,Obs}} = \frac{P_{\mathrm{F,M}\mapsto\mathrm{D}}}{[\mathrm{ARF}]_{\mathrm{T}}}\mathrm{FR}, \qquad [15]$$

$$k_{\mathrm{off,Obs}} = P_{\mathrm{D}\mapsto\mathrm{F,M}}\mathrm{FR}. \qquad [16]$$

We note that for the global fit the $k_{\mathrm{on}}$s in the transition probabilities were substituted with a single $k_{\mathrm{on}}$ ($k_{\mathrm{on,mic}}$, see Supporting note 1).

DRAFT
